# Extreme envelope plasticity drives temperature-dependent morphogenesis in the LPS-free bacterium *Sphingobium yanoikuyae*

**DOI:** 10.64898/2026.06.04.730141

**Authors:** Samantha Crahay, Henri Voedts, Nils Y. Meiresonne, Linus Wilhelm, Erika Hendrickx, Jean-François Collet, Katy Poncin

**Affiliations:** Unité de Recherche en Biologie des Microorganismes (URBM), Narilis, University of Namur, Namur, Belgium; de Duve Institute, Université catholique de Louvain, Brussels, Belgium; Bacterial Cell Biology, Instituto de Tecnologia Química e Biológica António Xavier, Universidade NOVA de Lisboa, Oeiras, Portugal; Center for Microscopy and Molecular Imaging (CMMI), Université Libre de Bruxelles, Gosselies, Belgium; Current affiliation: Biochemistry and Genetics of Microorganisms (BGM), Louvain Institute of Biomolecular Science and Technology (LIBST), Université catholique de Louvain, Louvain-la-Neuve, Belgium

## Abstract

Bacterial growth patterns are generally considered to be constrained and species-specific. Here, we show that *Sphingobium yanoikuyae* exhibits exceptional plasticity in envelope architecture and morphogenesis supported by a glycosphingolipid (GSL)-based outer membrane and an atypical peptidoglycan structure. At 30°C, cells expand asymmetrically via a rare bipolar envelope synthesis mode, whereas growth at 37°C triggers a transition toward longitudinal elongation accompanied by increased outer membrane vesiculation, indicating a reversible reprogramming of morphogenesis under host-relevant temperature. This switch is supported by rapid outer membrane dynamics, including high membrane fluidity and fast redistribution of envelope components. In addition to these pronounced morphological changes, peptidoglycan contains unusually short glycan strands and displays specific temperature-dependent remodeling suggesting that growth plasticity is not only governed by spatial regulation but also by changes in cell wall cross-linking pattern. Genetic analyses further identify essential roles for proteins involved in outer membrane–peptidoglycan and outer membrane–inner membrane coupling, as well as GSL transport systems, and core cell wall synthesis machinery, while revealing extensive redundancy in envelope remodeling enzymes. Together, these results establish *S. yanoikuyae* as a model for extreme envelope adaptability, where a highly fluid outer membrane and structurally unconventional peptidoglycan enable reversible transitions between distinct growth programs, potentially shaping environmental fitness and host-associated interactions.

## Introduction

In 1994, the team lead by Y. Koezuka discovered Agelesphins, bioactive molecules that are naturally produced by the marine sponges *Agelas mauritianus*^1^. These compounds have potent anti-tumor properties through their activation of invariant natural killer T (iNKT) cells^2^. Contrarily to classical antigens, they are not proteinaceous but are instead glycosphingolipids (GSL) composed of a ceramide backbone linked, in an α-conformation, by a single galacturonic acid sugar. For years, this remained a conundrum, as it was very unlikely that the human or mice immune system would have evolved to recognize marine sponge antigens. It was finally established that the organisms at the heart of this immune response were bacteria, mainly of the Sphingomonadales clade^2–4^ which belong to the α-proteobacteria class, a lineage of major evolutionary significance because the ancestors of mitochondria emerged from within this group^5^. A striking characteristic of Sphingomonadales is that despite being Gram-negatives, their outer membrane lacks lipopolysaccharides (LPS); instead, these molecules are functionally replaced by GSL. The GSL of *Sphingobium yanoikuyae* are remarkably similar to the α-Galactosylceramide (α-GalCer) molecules used in preclinical and clinical trials against various types of cancers^6^. It is therefore without surprise that purified *S. yanoikuyae* GSL have been shown to be activators of iNKT cells, NK cells, dendritic cells, and macrophages^7^. Strikingly, in a study published in 2020, *S. yanoikuyae* was detected in 36% of bone cancer samples but also as a commensal of healthy breast tissues^8^.

Despite their potent immunomodulatory properties, little is known about these bacteria as a whole. It remains unclear how GSL are produced and exported and how such a change to the outer membrane affects their physiochemical properties. Understanding bacterial growth and envelope biogenesis is essential, as these processes influence membrane composition, surface exposure of bioactive molecules, and adaptation to environmental or host-associated conditions. This question has been extensively addressed for α-proteobacteria, which display diverse growth strategies linked to lifestyle^9^. For example, *Agrobacterium tumefaciens* grows unipolarly, directing exopolysaccharide secretion to the old pole while generating progeny at the opposite pole, whereas *Caulobacter crescentus* elongates longitudinally but differentiates into motile and sessile cell types^9^. Given the unique GSL-rich outer membrane of *S. yanoikuyae*, characterizing its growth pattern and envelope organization is a prerequisite to understanding its physiology and host interactions.

To address these questions, we combined live-cell fluorescence microscopy, peptidoglycan labeling, biochemical analyses of cell wall architecture, and transposon sequencing to identify genes required for growth and envelope integrity. These experiments reveal that *S. yanoikuyae* grows *via* a rarely described “new end take-off” (NETO) mechanism and show how this asymmetric expansion is accommodated, defining the structural and genetic features that support bacterial growth. Strikingly, when cells are grown at 37°C, a condition that mimics host association, the growth pattern shifts toward predominantly longitudinal expansion, revealing a high degree of phenotypic plasticity in morphogenetic organization. Collectively, our findings provide a framework for understanding how an atypical, GSL-rich envelope shapes bacterial physiology across distinct environmental contexts and lays the groundwork for exploring potential host interactions.

## Results

### Microscopy reveals asymmetric cell division and cell length heterogeneity

*S. yanoikuyae* occupies a wide range of niches, including the rhizosphere of rice and rapeseed, where it promotes seed germination and root development^10,11^, as well as human-associated environments such as healthy breast tissues^8,12^, breast milk^13^, and bone cancers^8^. To investigate bacterial morphology in a context relevant to its ecological diversity, we first examined its growth in liquid cultures at different temperatures. Phase-contrast microscopy revealed marked morphological differences between cells grown at 30°C, representative of environmental conditions, and 37°C, representative of human-associated conditions. Although *S. yanoikuyae* displayed a broad distribution of cell lengths under both conditions, the proportion of longer cells was clearly increased at 37°C (**Fig. 1AB** and **Suppl. Fig. 1**). Despite this variability, septum placement - occasionally more than one per cell - was invariably asymmetric at both temperatures, indicating that *S. yanoikuyae* undergoes asymmetric cell division (**Fig. 1B**). Scanning electron microscopy revealed more subtle details such as the rare formation of Y-shaped poles is some cells grown at 30°C (top right panel, **Fig. 1C**), typically seen in polarly growing cells^14,15^, and the generation of outer membrane vesicles (OMVs) when grown at 37°C (bottom right panel, **Fig. 1C**), suggesting a reduced outer membrane stability and/or weaker outer membrane-peptidoglycan interactions^16^.

**Fig. 1.**
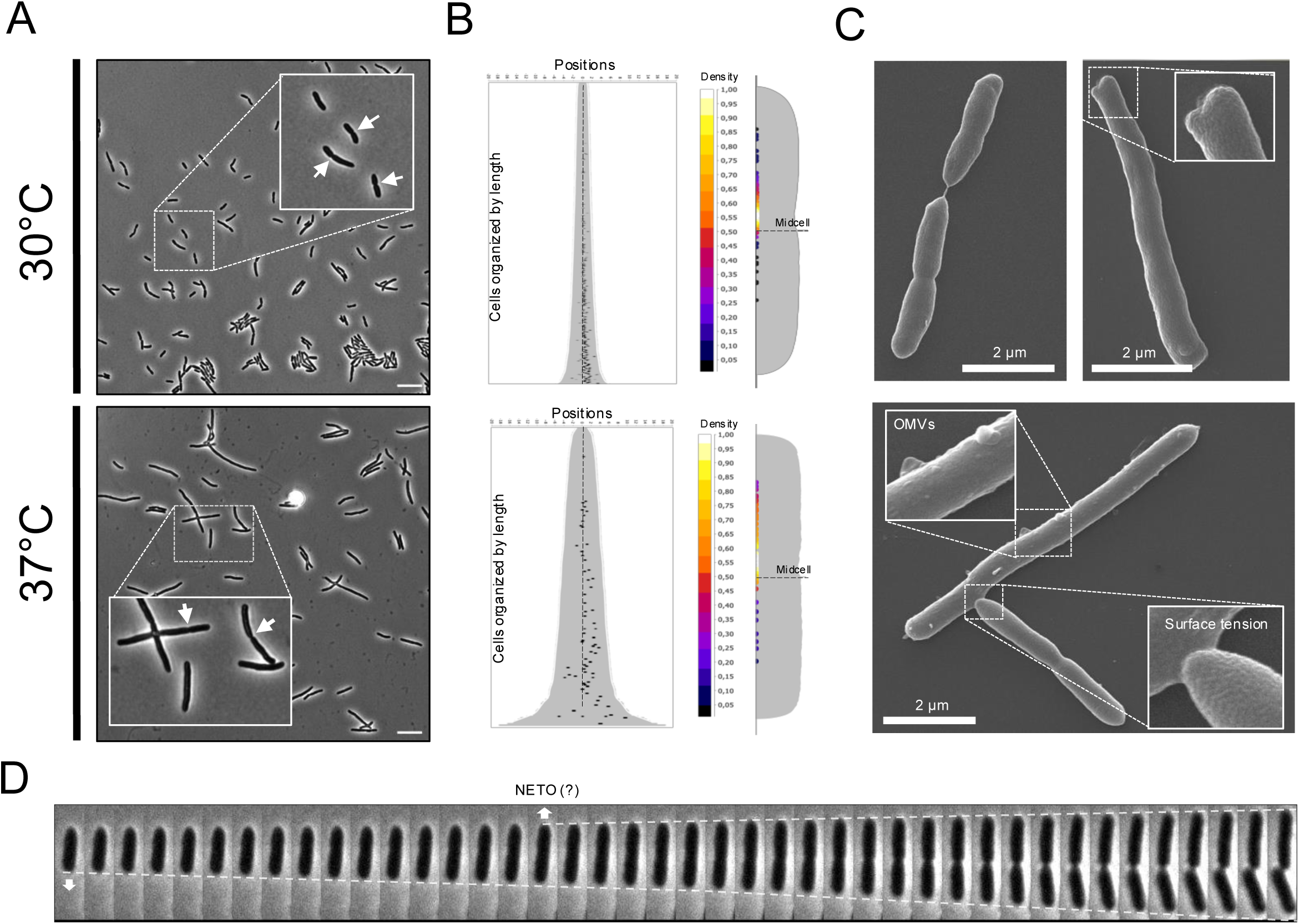
Microscopy images of *S. yanoikuyae* grown at 30°C and 37°C in liquid medium. **A.** Phase contrast microscopy. White arrows point to septa sites. Scale bars represent 10 μm. Violin plots representing cell length distributions can be found in Suppl. Fig.1. **B.** On the left, demographs representing septal positions mapped on cells organized by length (n = 479 and 209 cells, respectively). On the right, density map of septa on a representative bacterium (n = 195 and 66 septa, respectively). **C.** Scanning electron microscopy with dividing bacteria grown at 30°C (top left panel), aberrant Y-shape bacterium grown at 30°C (top right panel) and outer membrane vesicles (OMVs)-producing bacteria at 37°C (bottom panel). **D.** Time lapse images of a bacterium cultured on TSB pad at 30°C. Doted lines show growth evolution. White arrows represent apparent sites of growth. NETO stands for New End Take Off, when the second pole starts having visible apparent growth. Images were taken every 2.5 minutes.

Asymmetric growth patterns are not unusual in α-proteobacteria^9^. Indeed, *Hyphomicrobiales*, comprising the animal pathogen *Brucella abortus* and the plant pathogen *A. tumefaciens*, elongate unipolarly^17^. In contrast, members of the *Caulobacteraceae* family, such as *C. crescentus*, usually possess asymmetric polar appendages and grow through midcell elongation, with species-specific additional characteristics, such as growth directionality and extra disperse or polar growth zones^18^. Another growth pattern that can give rise to asymmetric daughter cells is the NETO mechanism, which for years was thought to be specific to eukaryotic organisms such as the fission yeast *Schizosaccharomyces* pombe^19^. It is now established that *Mycobacteriales*, monoderm bacteria including *Mycobacterium tuberculosis*^19^ and *Corynebacterium glutamicum*^20^ also use this growth pattern. Time lapse experiments were therefore performed directly on agar pad with *S. yanoikuyae.* At 30°C, the growth pattern was reminiscent of NETO growth, with one pole seemingly starting to grow before the second one takes off, leading to an increased cell elongation rate with time (**Fig. 1D**). Despite their ability to grow well in liquid culture at 37°C, growth on pad was impaired at this temperature, potentially due to the marked generation of OMVs (**Suppl. Fig. 2**). This was not unexpected as growth on petri dishes is also much arduous at 37°C compared to 30°C (about 100 h *vs* 30 h for colony forming units to appear).

### Fluorescent tagging uncovers temperature-dependent growth zones

To assess whether our observations reflect true NETO growth, we examined genetic markers associated with this growth pattern. In *Mycobacteriales*, a key marker of NETO is the DivIVA protein (also known as Wag31), which arbors a bipolar localization pattern with different intensities, as well as a pre-septal localization^20,21^. Though they are not homologous in sequence, DivIVA and the α-proteobacteria PopZ protein have been compared before, as they both function as polar scaffolding proteins, organizing cell-cycle and segregation machinery at bacterial cell poles^22^. In the unipolar growing *Hyphomicrobiales*, PopZ is found exclusively at the growing pole^23,24^, whereas it is unipolar or bipolar in the longitudinally growing *C. crescentus*, depending on its cell cycle progression^25^. Strikingly, in a *popZ::popZ-mChXL S. yanoikuyae* mutant, we observed an asymmetric bipolar localization of PopZ, as well as a pre-septal localization (**Fig. 2A**), reminiscent of what is observed for DivIVA in *Mycobacteriales*. Time lapse microscopy confirmed that the observed septal localization was the results of PopZ relocalization from the poles, rather than being inherited following division (**Fig. 2B**). Note that at 37°C, the bipolar localization remained but was accompanied by discreet widespread patches (**Fig. 2A**, bottom panel).

**Fig. 2.**
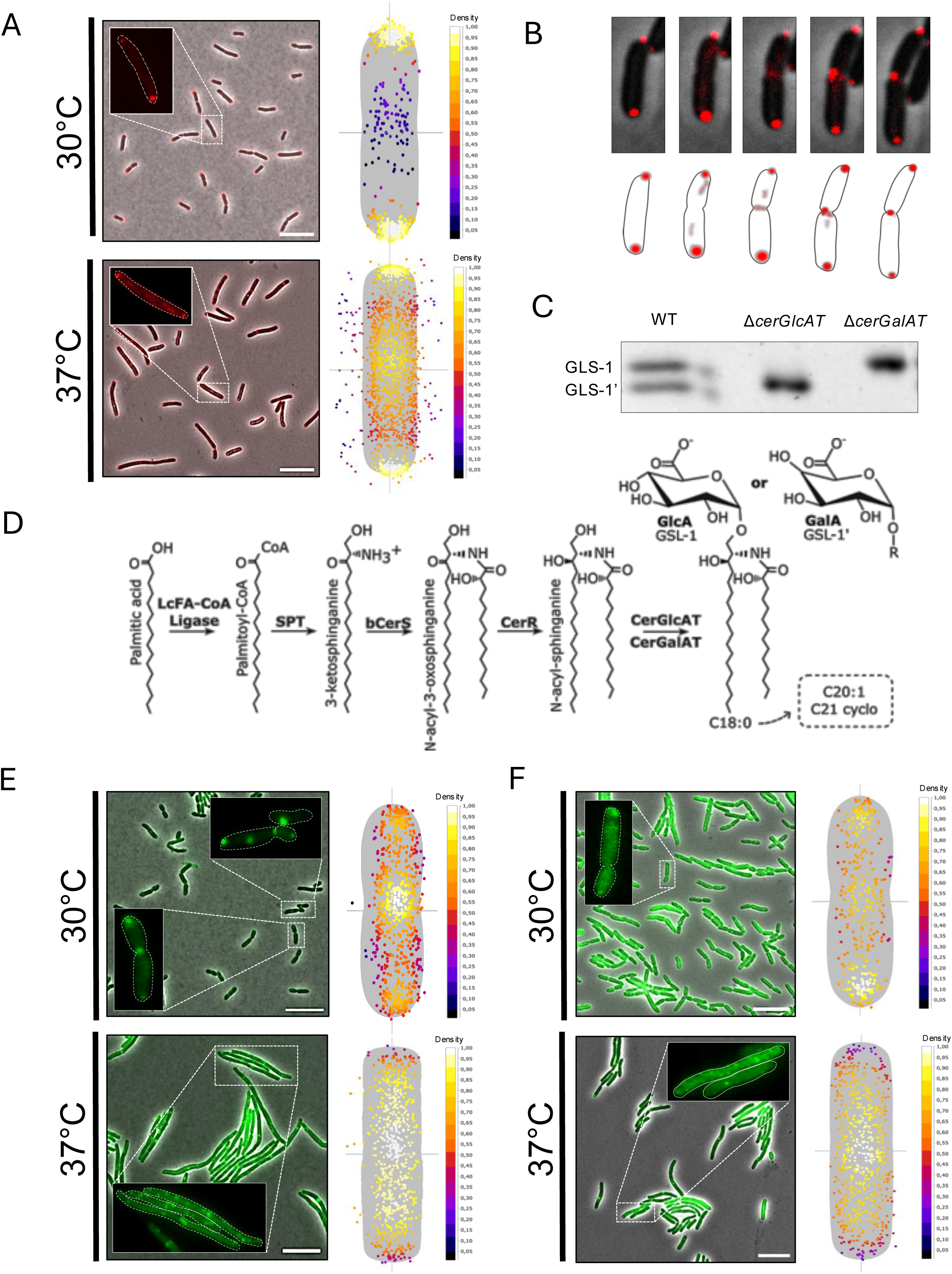
Growth zones determination through fluorescence tagging. **A.** PopZ-mChXL localization at 30°C and 37°C. On the left, representative microscopy images. Note the asymmetry in strength of the fluorescent signal between the two poles at 30°C. On the right, density maps of PopZ-mChXL foci on representative bacteria (n = 585 and 1726 foci). Scale bars represent 10 μm. **B.** Time lapse of the *popZ::popZ-mChXL* strain at 30°C. Images were taken every 5 minutes. **C.** Thin layer chromatography of glycosphingolipids (GSL) in the wild type, Δ*cerGlcAT* and Δ*cerGalAT* strains. **D.** Schematic representation of *de novo* GSL synthesis pathway from palmitic acid. Note that variations in the ceramide chain length and saturation status exist (dotted box). **E.** CerGlcAT-mNG localization at 30°C and 37°C. On the left, representative microscopy images. On the right, density maps of CerGlcAT-mNG foci on representative bacteria (n = 670 and 656 foci). Scale bars represent 10 μm. **F.** CerGalAT-mNG localization at 30°C and 37°C. On the left, representative microscopy images. On the right, density maps of CerGlcAT-mNG foci on representative bacteria (n = 345 and 631 foci). Scale bars represent 10 μm.

To determine whether our observations correspond to a true NETO pattern, defined as asymmetric bipolar growth, we planned to use the subcellular localization of proteins involved in GSL synthesis as a proxy for sites of active cell envelope growth. Indeed, in closely related unipolarly growing bacteria, the LPS synthesis machinery is tightly spatially regulated as to synchronize the synthesis of the outer membrane with the rest of the cell cycle^17^. Even though the two GSL coating the surface of *S. yanoikuyae* have long been described, the enzymes involved in their synthesis have yet only been inferred. Both GSL share the same lipid backbone upon which glucuronic acid or galacturonic acid are transferred to form GLS-1 and GLS-1’ respectively^26^. These additions of the terminal sugar moieties are catalyzed by terminal ceramide glycosyltransferase (CerGT) for which *S. yanoikuyae* encodes two; *cerGlcAT_1_* and *cerGlcAT_2_*. To infer their respective functions, we generated a markerless deletion mutant for each, extracted their lipids and visualized them by thin layer chromatography using a GLS specific stain. As expected, *S. yanoikuyae* had two distinct GLS that we attributed as being GLS-1 and GLS-1’ using previously published TLC^27^. The deletion of *cerGlcAT_1_* and *cerGlcAT_2_* resulted in the absence of GLS-1 and GLS-1’ respectively. CerGlcAT_1_ (thereafter CerGlcAT) and CerGlcAT_2_ (thereafter CerGalAT) are thus, in the tested conditions, specifically catalyzing the addition of GlcA and GalA on the sphingolipid, respectively. Next, to assess their localization, the two genes were fused to a fluorescent protein by allelic exchange. Strikingly, both enzymes revealed bipolar and pre-septal localization patterns at 30°C (**Fig. 2EF**). Nonetheless, slight variations were visible, such as sharper spotty signal with CerGlcAT-mNG compared to CerGalAT-mNG, and a propensity for the first to be found at the septum, while the other was preferentially found at the poles (**Fig. 2EF**). At 37°C, both enzymes had a stark delocalization pattern. Indeed, they were both excluded from the poles and were instead distributed alongside the cell body, with a preference for the midcell position (**Fig. 2EF**). Together, these localization patterns strongly support NETO growth at 30°C. Intriguingly, at 37°C there seems to be a more classical, *Escherichia coli*-like dispersed longitudinal growth.

### *S. yanoikuyae* displays high outer membrane fluidity

Historically, growth patterns have often been determined using fluorescent succinimidyl ester-based probes^17^. These molecules react with free primary amines of exposed molecules, *i.e.* with the N-terminus and lysine residues of cell surface proteins. Pulse-chase experiments therefore allow to follow the disappearance of the fluorescent probe at growing sites. In fully labeled unipolarly growing bacteria, the signal disappears from the new pole, as new proteins are incorporated along cell growth^17^. In contrast, in the laterally growing *Escherichia coli*, the signal becomes diluted in the sidewalls but remains static at the poles, where no growth occurs^17^. When a similar experiment was performed on *S. yanoikuyae*, we were unable to determine any clear growth pattern (**Suppl. Fig. 3)**. Considering bacterial GSL are known to have higher fluidity than LPS^28^, we hypothesized that outer membrane proteins might redistribute along the cell surface on timescales too fast for conventional succinimidyl ester-based labeling to reliably capture growth patterns. To test this, fluorescence recovery after photobleaching (FRAP) experiments were performed on labeled bacteria placed on plain PBS agar pads at 21°C, a condition with reduced fluidity and minimal growth. Images were taken every second following random pole photobleaching. Normalized FRAP curves appear to plateau at ∼50% of pre-bleach intensity within the first minute of imaging (**Fig. 3A**). However, both the initial slope and visual inspection of the bleached region indicate that full fluorescence recovery occurs usually within ∼10 seconds (**Fig. 3B**). The apparent plateau arises from partial bleaching of the reference region of interest in this highly fluid system, which underestimates the true recovery. These results indicate that *S. yanoikuyae* outer membrane proteins, embedded within the GSL layer, are actively moving along the bacterial surface, even in the absence of growth.

**Fig. 3.**
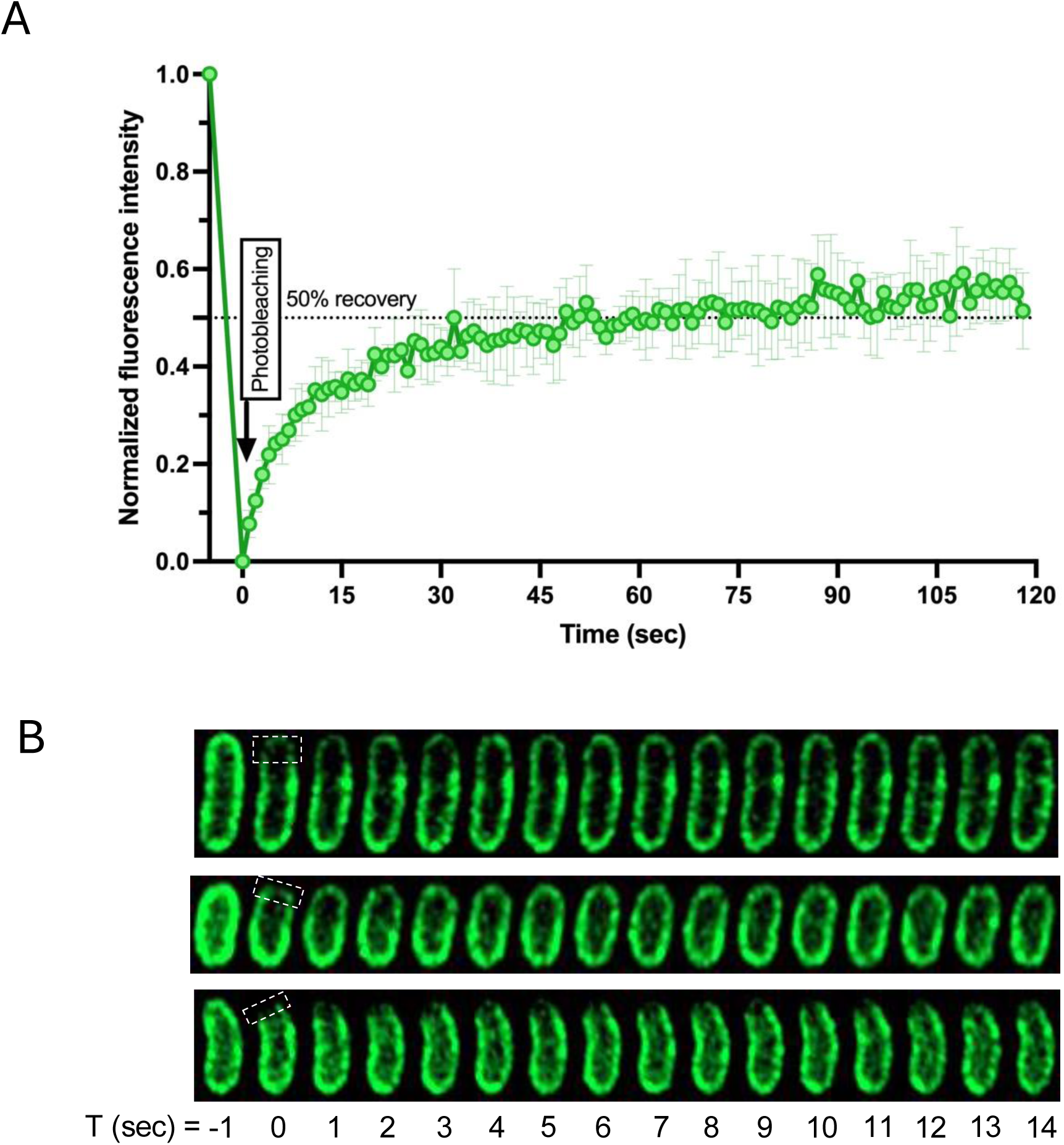
Fluorescence recovery after photobleaching experiments (FRAP). **A.** Normalized fluorescence intensity recovery over time, where 1 corresponds to pre-bleaching intensity fluorescence and 0 corresponds to the bleached frame. Mean values of three biological replicates (n = 32, 35, 33 bacteria). Error bars represent standard deviation from the mean. **B.** Time lapse images of three representative bacteria after photobleaching (T_0_). The white box represents the photobleached zone. The first two bacteria seemingly recover their full fluorescence within 10 seconds, whereas the third one takes longer.

### Uniform peptidoglycan synthesis supports growth and membrane remodeling

Peptidoglycan and outer membrane synthesis are tightly coordinated, as the outer membrane must remain physically linked to the underlying peptidoglycan layer to maintain envelope architecture, structural integrity, and resistance to environmental stresses^29^. In *S. yanoikuyae*, the occurrence of NETO growth coupled with rapid outer membrane reshuffling raises the question of how peptidoglycan synthesis accommodates such an unusual growth pattern, given that this rigid mesh-like structure has fundamentally different properties compared to the surrounding lipid layers. In *Mycobacteriales*, peptidoglycan insertion occurs predominantly at the cell poles, and follows a NETO-like biphasic pattern that parallels polar envelope expansion^30^. To check whether this is also true for *S. yanoikuyae*, the fluorescent D-amino acid (FDAA) HADA was used to label its peptidoglycan, as this molecule highlights transpeptidation reactions and allows direct observation of peptidoglycan crosslinking activities^31^. Surprisingly, evenly widespread labeling of the lateral and polar peptidoglycan was visible very shortly after contact with HADA, suggesting unusual high activity of periplasmic transpeptidases^32^ in a homogenous widespread manner (**Fig. 4A**). To make sure this result was not due to insufficient washing of excess probes, a rotor-fluorogenic DAA (rfDAA) named rf470DL was employed. Indeed, rfDAAs enable peptidoglycan labeling without a need for washes, as they emit fluorescence only in congested environment, *i.e.* upon their transpeptidation into the peptidoglycan mesh^33^. Mirroring the HADA labeling pattern, rf470DL incorporation revealed a fast and uniform signal across the cell surface, both at 30°C and 37°C (**Fig. 4A**). To gain further insight into the rapid and spatially widespread transpeptidase-mediated incorporation, we compared the HADA-labeling signals obtained after 5 min in the lateral walls and septa of pre-divisional bacteria grown at 30°C and 37°C. At both temperatures, periplasmic transpeptidase activity was higher at the septum than along the lateral walls. However, this difference was markedly reduced at 37°C, with a septum-to-side-wall ratio of 1.16 ± 0.18 compared with 1.43 ± 0.13 at 30°C (**Suppl. Fig. 4**). These results suggest that, at the host-associated temperature, periplasmic transpeptidase activity is relatively redistributed toward elongation rather than division.

**Fig. 4.**
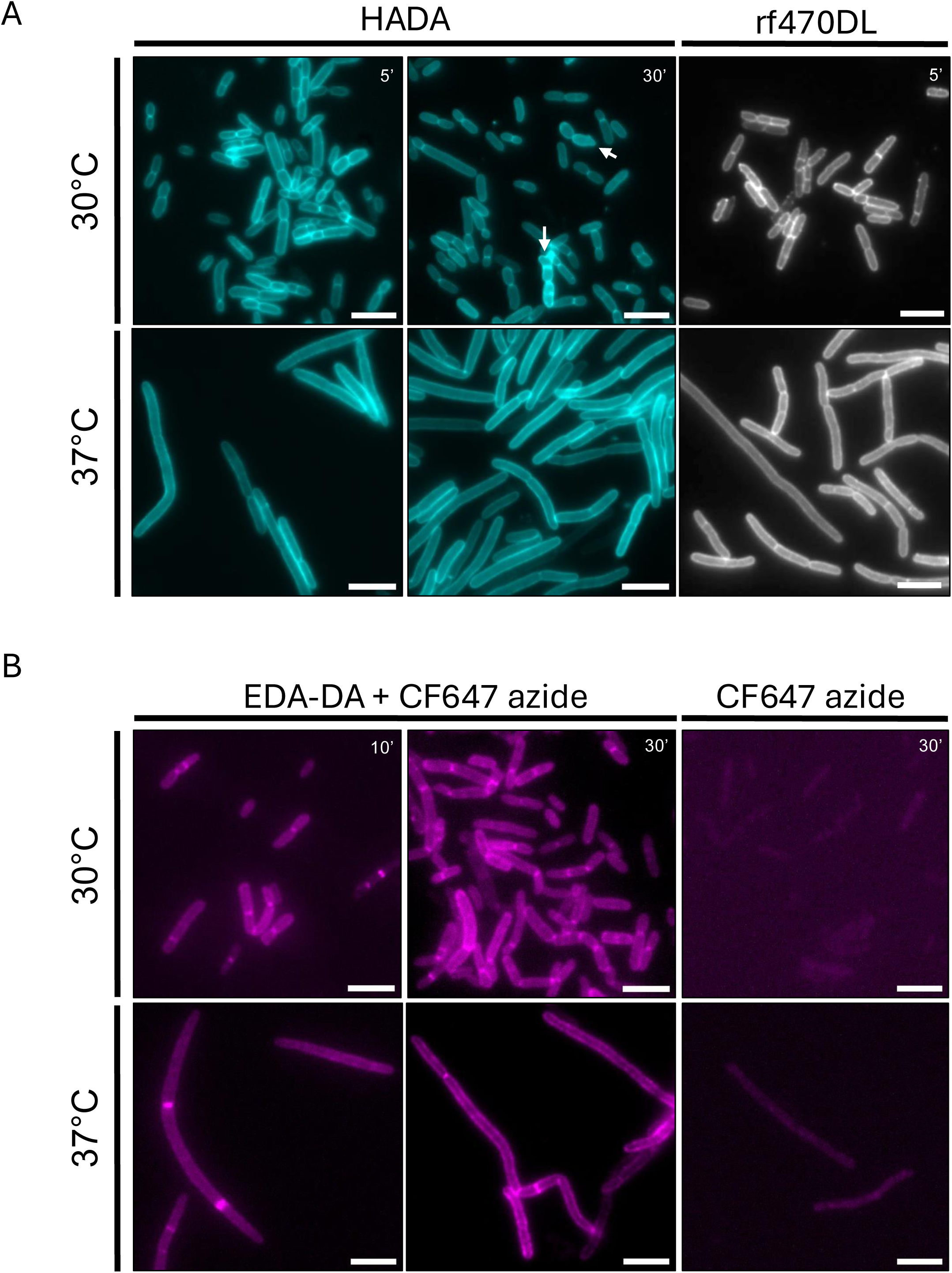
Peptidoglycan labeling by fluorescent probes. **A.** Peptidoglycan labeling through the activity of periplasmic LD-transpeptidases and DD-transpeptidases. The HADA probe is a fluorescent D-amino acid, whereas rf470DL is a rotor-fluorogenic D-amino acid, which requires crosslinking to the peptidoglycan to emit fluorescence. Scale bars represent 5 μm. White arrows point to bacteria undergoing Y-shaped polar growth art 30°C. Note that when labeling was prolonged for 30 min (∼33% of doubling time), Y-shaped bacteria were found more often than in unlabeled bacteria (∼5% compared to <1%). **B.** *De novo* peptidoglycan labeling. The clickable D-amino acid dipeptide EDA-DA was incorporated for 10 or 30 minutes within nascent peptidoglycan through the cytoplasmic pathway, then labeling was achieved with the rotor-fluorogenic CalFluor647 (CF647) azide probe, mixed with the outer membrane permeabilizer Polymyxin B nonapeptide hydrochloride (PMBN). The negative control (right panels) corresponds to the same protocol realized without EDA-DA incorporation. Scale bars represent 5 μm.

Since both FDAA and rfDAA reflect the activity of periplasmic transpeptidases, but not *de novo* peptidoglycan synthesis, clickable D-amino acid dipeptides (DAADs) were used next. EDA-DA is a D-alanine-D-alanine analog that gets incorporated directly into growing peptidoglycan peptide chains through the cytoplasmic MurF enzyme^34^. After allowing 10 and 30 minutes of EDA-DA incorporation in rich medium, *S. yanoikuyae* cells were washed and labeled with clickable rotor-fluorogenic CalFluor 647 Azide probe^35^ supplemented with the outer membrane permeabilizer Polymyxin B Nonapeptide hydrochloride (PMBN). As expected, a stronger signal was observed at the septum than at the side walls. Nevertheless, the signal along the side walls remained homogeneous at both 30°C and 37°C (**Fig. 4B**).

### A specialized peptidoglycan architecture supports envelope remodeling

As indicated above, the strikingly fast incorporation of HADA suggests that peptidoglycan remodeling must be very active. To better understand how peptidoglycan synthesis can accommodate enhanced outer membrane fluidity, we performed sacculus compositional analysis and found that *S. yanoikuyae* displays a highly atypical cell wall architecture (**Fig. 5**). Glycan strands were markedly short, with an average length of only ∼ 6-7 NAG-NAM repeats, suggesting extensive turnover and fragmentation of the glycan backbone, a rare featured shared with the flexible bacterium *Helicobacter pylori*^36^. Overall, only limited compositional differences were observed between peptidoglycan purified from bacteria grown at 30°C and 37°C, but several features pointed toward enhanced remodeling at the lower temperature through L,D-transpeptidase activity. The peptidoglycan network was dominated by classical 4→3 cross-links, although 3→3 cross-links were also detectable and were predominantly observed in bacteria grown at 30°C, contrasting with several polar-growing bacteria in which L,D-transpeptidase-mediated crosslinking contributes substantially to cell wall structure^30,37^. Consistent with increased L,D-transpeptidase activity, partial replacement of the terminal D-Ala⁴ residue by glycine was observed at 30°C but was not detectable at 37°C. This type of modification is known to lead to altered cross-linking patterns and reduced peptidoglycan rigidity in other bacteria^38^. Despite this, in *S. yanoikuyae*, total crosslinks were higher at 30°C than at 37°C (42.7% *vs* 34.1%), an increase that was largely attributable to the additional 3→3 cross-links detected at 30 °C. Tetramers of four cross-linked subunits were also detectable only at 30 °C. Together, these features point to a mechanically permissive and highly modulable peptidoglycan mesh, consistent with the need to accommodate high fluidity of the outer membrane components during asymmetric NETO growth or longitudinal elongation, a mechanism that differs from what is observed in *Mycobacteriales*^30^.

**Fig. 5.**
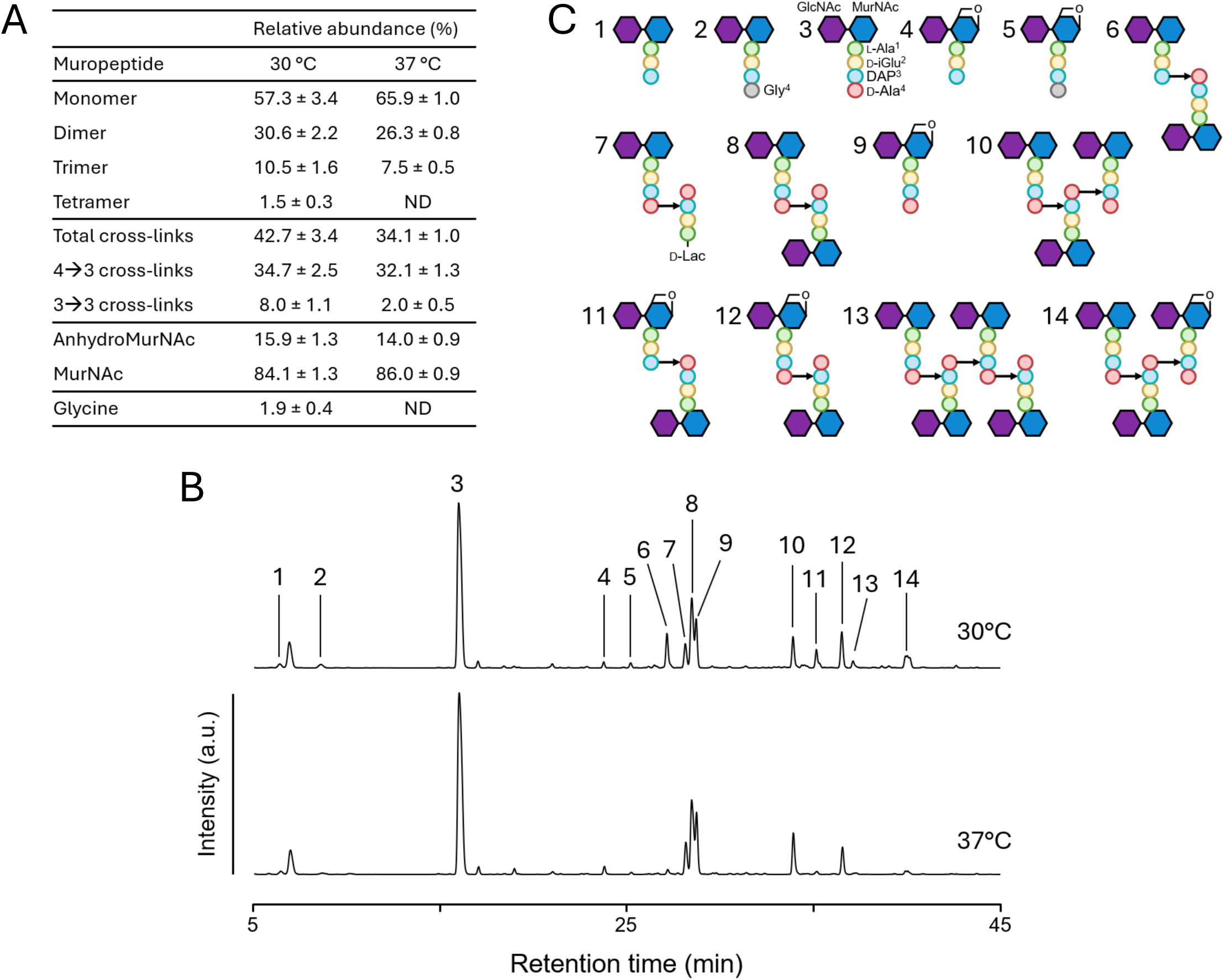
*S. yanoikuyae* peptidoglycan analysis. **A.** Peptidoglycan composition obtained from exponentially growing liquid cultures of *S. yanoikuyae* at 30°C and 37°C in TSB. Peptidoglycan was purified, digested into muropeptides, and analyzed by LC-MS. Values for each muropeptides are indicated in % of the total area. Considering that each glycan strand has one AnhydroMurNAc end, the average glycan chain length is ∼6-7 subunits. Data represent the mean ± standard deviation (n = 3 biological replicates). ND, not detected. **B.** Representative LC-MS profiles of muropeptides extracted from *S. yanoikuyae* grown at 30°C and 37°C in TSB. a.u., arbitrary unit. **C.** Structure of the muropeptides annotated in panel B. The peaks 4, 5, 9, 11, 12, and 14 correspond to muropeptides containing an AnhydroMurNAc.

### Tn-seq highlights cell wall, membrane, and cytoskeletal contributors to growth

To get better insights into the different actors needed for this peculiar growth plasticity, transposon libraries followed by deep sequencing (Tn-seq) was performed on *S. yanoikuyae*. About 1 million mutants were generated at 30°C with a mini-Tn5, for a genome of 5.5 Mb. Of these, 371,779 unique insertion sites were mapped on the reference genome. Data were then analyzed using a sliding window method coupled to the annotation-dependent TniF to determine essentiality (**Suppl. Table 1**)^39^. Note that the same experiment was not attempted at 37°C, since, as mentioned before, CFU formation is sub-optimal at this temperature.

Tn-seq revealed that envelope integrity and membrane organization constitute the primary genetic constraints underlying the bipolar growth program of *S. yanoikuyae*. As expected, several genes associated with GSL metabolism^40^ and transport^41^ were required for survival (**Fig. 6A , Suppl. Table 1**). Based on annotation and essentiality, we also identified a gene which could be responsible for the desaturase activity leading to variation in the ceramide chain depicted in **Fig. 2D**. Core peptidoglycan precursor synthesis genes remained essential, whereas multiple peptidoglycan remodeling enzymes and elongation-associated penicillin-binding proteins displayed extensive redundancy (**Fig. 6A**), consistent with a morphogenetic system that relies less on specialized cell wall chemistries than on spatial regulation of envelope expansion. In contrast, genes involved in outer membrane biogenesis and maintenance, including those of the Bam, Lol, and Tol-Pal systems, were all essential (**Fig. 6A**), highlighting the central importance of outer membrane assembly during growth at 30°C. Notably, the outer membrane-peptidoglycan coupling protein OmpA exhibited a strong essentiality signature (**Fig. 6A**), suggesting that mechanical linkage between envelope layers is critical for maintaining polarized growth. Among the most unexpected findings was the strong fitness contribution of a Mic60/mitofilin-like membrane organizer (**Fig. 6A**). A recent study identified bacterial Mic60 homologs in α-proteobacteria and demonstrated their association with membrane organization complexes through interactions with the outer membrane assembly factor BamA, consistent with a conserved role in envelope architecture^42^.

**Fig. 6.**
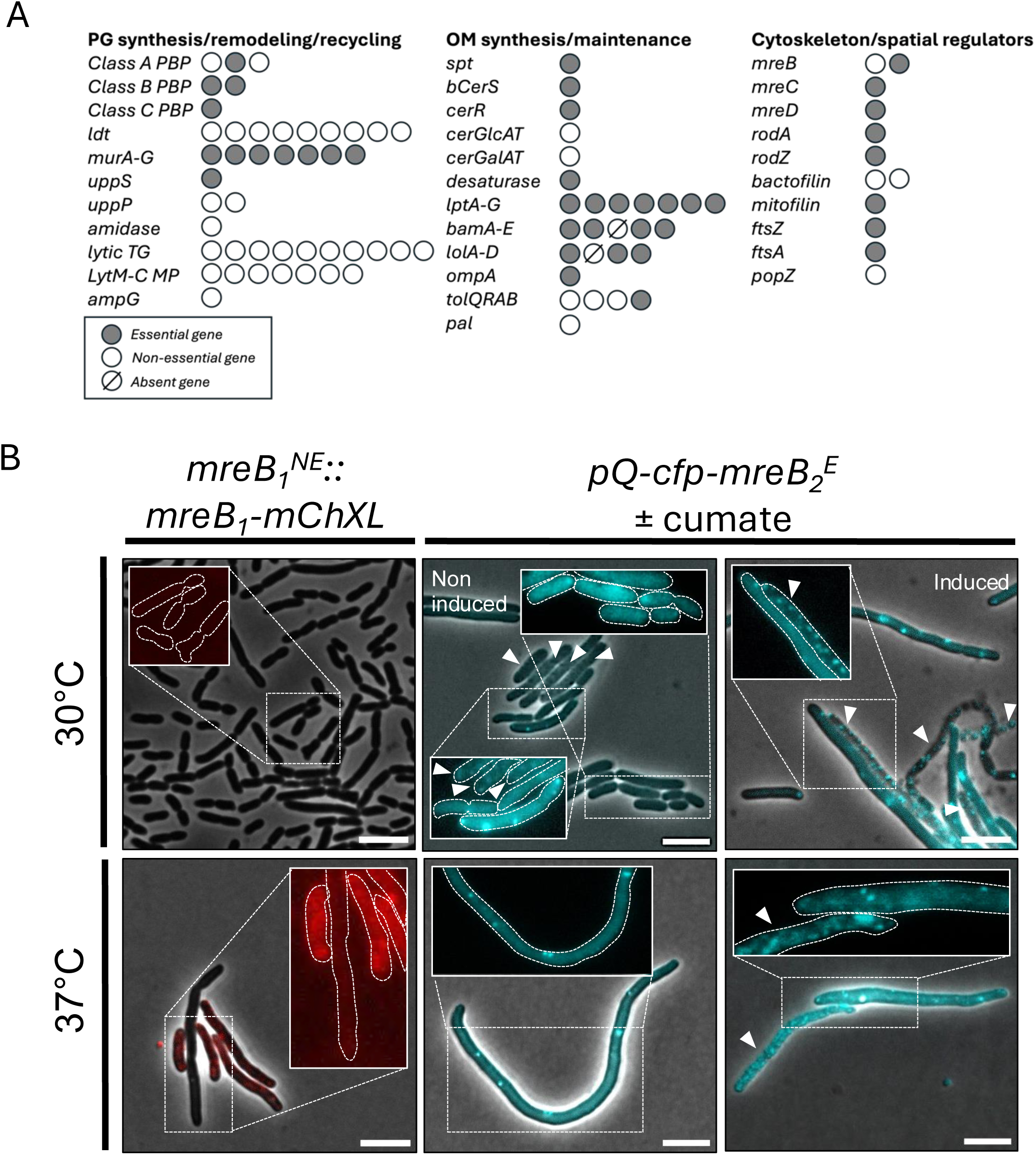
Cell wall, membrane, and cytoskeletal contributors to growth. **A.** Tn-seq data revealing essential genes on all three gene categories. The number of circles corresponds to the number of correspond genes annotated under the same name or with the same predicted function. *PBP* stans for *Penicillin Binding Protein, ldt* stands for *LD-transpeptidase, TG* stands for *transglycosylase, LytM-C MP* stands for *LytM (M23)-containing metallopeptidases*. Note that *lpt* genes are classified as involved in GSL transport, as supported by Uchendu *et al*, 2026. Details for each gene depicted here can be found in Suppl. Table 1. **B.** Microscopy images of the two MreB paralogs fused to fluorescent proteins. The non-essential (^NE^) *mreB_1_* was fused to a *mChXL* sequence through allelic replacement, while the essential (^E^) *mreB_2_* was fused to a *cfp* sequence on a cumate-inducible plasmid. White arrows point to bacteria undergoing lysis. Scale bars represent 5 μm.

Another striking finding was that despite their duplication, the two *mreB* genes had non-redundant functions, as only one of them was essential at 30°C (**Fig. 6A**). To get better insight into their respective roles, both genes were tagged with a fluorescent fusion; the non-essential *mreB_1_* directly by gDNA allelic replacement, and the essential *mreB_2_ via* a cumate-inducible plasmid. Remarkably, MreB_1_-mChXL displayed a fluorescent signal solely at 37°C, while CFP-MreB_2_ led to bacterial lysis, independently from its induction status (see white arrows on **Fig. 6B**). This unexpected divergence in essentiality and inducibility between the two MreB paralogs reveals strong functional specialization within what is typically considered a redundant elongasome system.

Together, these data indicate that morphogenesis during growth is strongly dependent on membrane organization and envelope connectivity rather than exclusively on dedicated peptidoglycan elongation systems.

## Discussion

Bacteria are exquisitely adapted to their environments, including through their growth mode strategies. *Sphingomonadales* are among the most ubiquitous bacteria^43^ and, despite extensive characterization of growth patterns within α-proteobacteria, their own growth dynamics remain poorly understood^9^. A striking characteristic of *Sphingomonadales* is the complete absence of LPS at their outer membrane. Indeed, their interface with the external environment is composed of GSL rather than LPS, conferring them unexpected permeability properties^43^ as well as specific cytotoxic activities^7^. Some species are even described to be able to perform phagocytic-like engulfment of complex sugars^44^. Here, we set out to study the growth pattern of *Sphingobium yanoikuyae*, as this bacterium is associated with marine environment^45^, contaminated soil^46^, roots^10^, but also human samples^12^, including bone cancers^8^.

The present study reveals that *S. yanoikuyae* possesses an unusually dynamic and plastic cell envelope organization that supports distinct modes of bacterial growth depending on environmental context. We found that *S. yanoikuyae* displays a NETO-like asymmetric growth of its outer membrane under environmental conditions, but undergoes substantial morphogenetic remodeling at 37°C, a temperature consistent with host association. This is possible through coupling to an unconventional peptidoglycan architecture, enabling remarkable flexibility in envelope organization and growth behavior. Indeed, the apparent presence of only ∼6-7 disaccharide units per strand (**Fig. 5**) would imply a highly truncated or extensively remodeled sacculus, which could significantly affect wall elasticity and porosity. Such shortened glycan chains are typically associated with either high turnover rates by autolysins and lytic transglycosylases or specialized structural adaptations where mechanical stability is achieved more through cross-link density than chain length^47,48^. Our data are in agreement with this, as we found extensive and rapid labeling of the peptidoglycan with the HADA probe (**Fig. 4A**), and a high level of redundancy within LDT, LytM-containing metallopeptidases and lytic transglycosylases (**Fig. 6A**). Importantly, the essentiality of both *ompA* and *mitofilin* (**Fig. 6A**) suggests that tethering of the outer membrane to the peptidoglycan^49^ and the inner membrane^42^, respectively, might be a key factor in allowing this high morphologic plasticity.

While multiple MreB homologs have been described in some bacterial species^50,51^, clear functional specialization associated with different morphogenetic programs remains relatively uncommon. Here, one paralog appears essential during NETO-like growth at 30°C, whereas the second is specifically expressed at 37°C, where cells undergo longitudinal elongation (**Fig. 6B**). This suggests that the two proteins may organize distinct elongasome activities or patterns of peptidoglycan insertion depending on environmental conditions. Interestingly, despite these major differences in growth pattern and presumed cytoskeletal organization, the overall peptidoglycan structure appears to remain largely similar at both temperatures (**Fig. 5**), except for the additional L,D-transpeptidase activity detected at 30 °C. This indicates that the bacterium may achieve different morphogenetic outcomes without substantially altering the biochemical composition of its cell wall, possibly by changing the spatial regulation or dynamics of peptidoglycan synthesis instead. Such a mechanism would support the idea that MreB paralogs can differentially control cell elongation programs while maintaining a conserved envelope architecture. One tempting hypothesis is that MreB_2_ regulates NETO growth while MreB_1_ regulates longitudinal elongation. Future studies should clarify if this is true by looking at the phenotypes of deletion/depletion strains.

The outer membrane of *S. yanoikuyae* is not composed of solely GSL-1 (α-GlcACer) but also of GSL-1’ (α-GalACer), with a mix of saturated and unsaturated ceramide acyl chains^2^ (**Fig. 2D**). This bears the question of their respective distribution and functions in both host-associated and environmental context, considering they appear to have different immunological^2^ and physiological^52^ properties. First, we established which transferase enzyme was responsible for which sugar grafting (**Fig. 2C**) and confirmed the predicted specificity of CerGlcAT for glucuronic acid^53^. We found that, despite being very similar, the two enzymes had slightly different localization patterns at 30°C, with CerGlcAT being mainly found at the pre-septal site (**Fig. 2E**), while CerGalAT tended to be localized preferentially at the poles and more diffused (**Fig. 2F**). At 37°C, both enzymes had a widespread spotty localization with exclusion form the poles (**Fig. 2EF**). Considering the extreme membrane fluidity we found at room temperature (**Fig. 3**), these different localization patterns should be considered in the context of direct contact sites. For environmental and plant-associated growth, polar outer membrane synthesis^16^ seems evolutional interesting, since both *A. tumefaciens* and rhizobia take advantage of this property to adhere polarly to their targets^54,55^. *Sphingomonadales* being producers of exopolysaccharides^56^, this could be a way for the bacteria to promote tip adherence, potentially through rosette formation^57^. In this context, it would therefore be interesting to investigate whether GSL can serve as lipid anchors to exopolysaccharides, similarly to LPS^58^. Transient condensed polar GSL-1’ could also serve as a signal for plant adhesion, as GalA – contrarily to GlcA – it is the main building block of pectin, a major component of the plant cell wall^59^.

The shift from polar growth to a more conventional longitudinal growth pattern at 37°C might also be linked to adhesion. Indeed, physiological studies comparing LPS to GSL revealed that GSL vesicles have stronger surface adherence properties^60,61^, which we also observed by transmission electron microscopy (**Fig. 1C**). At this host-associated temperature, one striking feature of *S. yanoikuyae* is its elongated phenotype (**Fig. 1A**). This is reminiscent of uropathogenic *E. coli* (UPEC) that filament in a SOS-independent manner to limit phagocytosis by macrophages^62^. It was also found that a delay in the phagocytosis of filamentous *L. pneumophila* was due to macrophages trying to reorient the bacteria to find the pole^63^. Importantly, *S. yanoikuyae*’s GSLs have previously been shown to promote polymorphonuclear leukocyte phagocytosis and lysosomal fusion while eliciting limited superoxide production^64^. In this context, the transition toward elongated morphologies and increased OMVs production at 37°C may reflect a specialized host-interaction state rather than a generic stress response. OMVs could contribute to the redistribution or release of immunologically active envelope components, potentially modulating interactions with phagocytic cells. Although these hypotheses remain speculative, the coordinated changes in morphology, membrane organization, and vesiculation observed at host-associated temperatures strongly suggest that *S. yanoikuyae* actively remodels its envelope architecture in response to environmental transitions.

Together, our findings support a model in which *S. yanoikuyae* employs a flexible envelope architecture capable of supporting multiple morphogenetic programs through dynamic reorganization of membrane synthesis, cytoskeletal activity, and peptidoglycan remodeling. More broadly, our study suggests that GSL-rich bacterial envelopes may represent an alternative strategy for outer membrane organization, combining structural stability with exceptional adaptability. Given that bacterial GSLs have also been associated with immunomodulatory and anti-tumor activities, it raises intriguing questions regarding their interactions with host cells and their potential role in cancer-associated environments. Understanding how these atypical envelopes influence microbial physiology and host interactions will provide important insight into the diversity of bacterial cell biology beyond classical model systems.

## Material and Methods

### Bacterial strains and media

*E. coli* strains DH10B (Thermo Fisher Scientific) and MFDpir^65^ were grown in lysogeny broth (LB) at 37°C. The type strain *S. yanoikuyae* DSM 7462 (equivalent of ATCC 51230; obtained from the Leibniz Institute DSMZ-German Collection of Microorganisms) and its derivatives were grown in TSB-rich medium (3% Bacto, tryptic soy broth), except if otherwise stated. Plating of *S. yanoikuyae* was always performed at 30°C, and liquid cultures were either done at 30°C or 37°C. Antibiotics were used at the following concentrations: kanamycin, 50 μg mL^−1^ for *E. coli* and 30 μg mL^−1^ for *S. yanoikuyae*; diaminopimelic acid (DAP), 0.3 mM; streptomycin 150 μg mL^−1^; polymyxin B 1 μg mL^−1^; tetracycline 10 μg mL^−1^.When required, cumate was used at 50 μg mL^−1^.

### Plasmid constructions and cloning procedures

*S. yanoikuyae* deletion strains and fluorescent fusion strains were constructed by allelic exchange, via pAK405 vectors (J. Vorholt, Addgene plasmid #37114) carrying a kanamycin resistance cassette and a streptomycin sensitivity cassette^66^. The upstream and downstream regions around the gDNA site of interest (most of the gene for deletion, and specific N- or C-terminal codons for fusions) were about 800 bp long and ligated by overlap PCR before being inserted into the pAK405 vector into SmaI restriction sites. We performed conjugation with the *E. coli* MFDpir strains carrying the plasmids and *S. yanoikuyae*, from colonies recovered from solid agar plates and spotted in 100 μl drops on TSB supplemented with DAP. After overnight incubation at room temperature, bacteria were spread on TSB plates supplemented with kanamycin and polymyxin B. After incubation at 30°C, 5-6 isolated *S. yanoikuyae* colonies were recovered with a loop and spread on TSB plates supplemented by streptomycin for counterselection to allow plasmid curing, then incubated at 30°C until isolated colonies appearance. To confirm gene deletion or fusion, PCRs were performed on colonies that were kanamycin-negative and streptomycin-resistant with primers targeting sequences upstream and downstream of the plasmid-containing sequences.

The pQ*-cfp-mreB_2_* inducible plasmid was generated by inserting a PCR containing the *mreB_2_* gene sequence into the pQC plasmid (J. Vorholt, Addgene plasmid #48100)^67^ using the XhoI and SpeI restriction sites. It was then transferred to *S. yanoikuyae* by conjugation with *E. coli* MFDpir as described above, and the final strain was selected on TSB plates supplemented with tetracycline and streptomycin.

For focal plan determination of FRAP experiments (see below), a stable constitutively fluorescent “Red” *S. yanoikuyae* strain was generated. Briefly, we used the allelic exchange method described above to insert a long PCR product (containing the following sub-parts: [upstream region]-[p ^68^]-[RBS]-[*mCherryXL*]-[downstream region]) into a pAK405 plasmid. The targeted genomic region was between the forward-facing genes with the ORFs LAHMBCNB_01869 and LAHMBCNB_01870. Note that the *mCherryXL* sequence^69^ was custom-made to be codon-adapted to *S. yanoikuyae* and ordered as GBlock (IDT).

Primers, plasmids and ORFs of the studied genes are listed in **Suppl. Table 2**.

### Probe-based peptidoglycan labeling

Exponential phase cultures of *S. yanoikuyae* were labelled with 500 μM HADA (Tocris Bioscience) for 5 or 30 minutes in 50-100 μl final volume, then cells were fixed by adding 1 mL of 70% ice cold ethanol for 5 minutes, followed by three washes in PBS before microscopy. Since the rotor-fluorogenic probe rf470DL does not enter readily in gram-negative bacteria^33^, including *S. yanoikuyae*, we set up a new protocol to circumvent this issue without needing to mutate the bacterial strain. Bacterial pellet was first washed once with PBS, then resuspended in 1 mM rf470DL (Tocris Bioscience) supplemented with 500 μg mL^−1^ of the outer membrane permeabilizer PMBN (Merck) in PBS, in 50-100 μl final volume. Cells were then fixed by adding 1 mL of 70% ice cold ethanol, followed by two washes in PBS before microscopy.

For EDA-DA labeling, cells were incubated with 2.5 mM EDA-DA (Tocris Bioscience) for 10 or 30 minutes, before being washed once in PBS, then labeled for 30 minutes with 40 μM of the rotor-fluorogenic clickable probe CalFluor 647^70^ azide (Vector Laboratories) using a BTTAA-based commercially available CuAAC Cell Reaction Kit (Jena Bioscience) according to manufacturer’s instructions, but with the addition of 500 μg mL^−1^ to the reaction buffer, in 50-100 μl final volume. Cells were then fixed by adding 1 mL of 70% ice cold ethanol, followed by two washes in PBS before microscopy.

### Epifluorescence microscopy and image analysis

Images were taken with an inverted microscope Nikon Eclipse Ti2 equipped with a phase-contrast objective Plan Apo λ DM100XK 1.45/0.13 pH3 and a Hamamatsu C13440-20CU ORCA-FLASH 4.0 camera (Hamamatsu). Bacteria in culture (2 μL) were observed with phase contrast on PBS-agarose (1%) pads or on TSB agar pads for time lapse experiments. Pictures were encoded with NIS-element software and analyzed with the plug-in MicrobeJ^71^ in ImageJ/Fiji^72^. For demographs, septum localizations were determined using the constriction feature detection parameters within the bacterial detection options. Fluorescent septa were localized using the Point tool within Maxima options. Septa or fluorescent spots patterns were visualized using the Subcellular Localization tool withing the Result interface. Cells were oriented such that the pole closest to a septum would be pole 1 (top) and the one furthest from it was pole 2 (bottom), and cells were aligned at the midcell.

Ratios of septal over lateral fluorescence from Suppl. Fig. 4 were obtained by measuring and dividing mean fluorescent signal of septal and lateral fluorescence signal of a single cell in ImageJ^72^. Regions of interests were drawn manually over septal signals and their four directly adjacent lateral signals. Regions where cells touch with other cells were excluded to prevent measuring double fluorescence signal. At least 50 individual cells of each group were analysed.

### Scanning electron microscopy (SEM)

Exponential phase bacteria (3 mL) were fixed for 1h at 160 rpm and at room temperature with 5% glutaraldehyde directly added to their growth medium. They were then pelleted at 4000 G and washed in 0.1 M cacodylate buffer before being resuspend 1 mL 0.1 M cacodylate buffer and stored at 4°C until further processing. Cells were deposited overnight onto poly-L-lysine-coated coverslips (1 mg/mL). Samples were then washed three times for 5 min each in 0.1 M cacodylate buffer and post-fixed with 2% osmium tetroxide in 0.1 M cacodylate buffer for 1h. After three additional 5 min washes in 0.1 M cacodylate buffer, samples were dehydrated through a graded acetone series (50%, 70%, 95%, and 3x100%) for 15 min at each step. Critical point drying was performed using a EM CPD030 system (Leica Microsystems), allowing the substitution of acetone with liquid CO₂ prior to vaporization. Samples were subsequently sputter-coated with a 20 nm platinum layer using a EM MED020 metallizer (Leica Microsystems). Imaging was carried out with an ESEM Quanta F200 (Thermofisher Scientific) scanning electron microscope equipped with an Everhart–Thornley secondary electron detector.

### Fluorescence recovery after photobleaching (FRAP)

To allow labelling of the outer membrane proteins of *S. yanoikuyae*, an alternative growth medium had to be used to limit exopolysaccharides production at its surface. The “Red” strain of *S. yanoikuyae* (see above) was first inoculated on TSB agar plates and incubated at 30°C for about 30 hours, before isolated colonies were recovered with a full loop and transferred in 1 mL of M2 minimal medium^73^ supplemented with 0.05% glucose. Bacteria were then separated by pipetting and left to sediment for 10 seconds. The non-sedimented bacteria in suspension were then transferred into 5 mL culture of M2G and incubated overnight at 37°C. Note that under these specific conditions, bacteria barely elongate. The next morning, bacteria were washed in 50% PBS, then 75% PBS for osmotic pressure stabilization, before being labelled during 15 minutes at room temperature with AlexaFluor™ 488 NHS Ester (Invitrogen) at a final concentration of 10 μg mL^−1^ in PBS. Bacteria were then washed three times in PBS and observed with a Zeiss LSM900 Airyscan2, objective 63x (PL-Apo, NA 1.4 oil) equipped with a Axiocam 305 mono camera (Zeiss). To check viability and find focal plan, bacteria were observed with the mChXL spectra, then FRAP was performed with the following parameter: crop area of 1.7x, intensity laser at 100%, time series of 2 minutes with intervals of 1 second, smart autofocus activated. Small square zones of photobleaching were defined for random bacterial poles and FRAP analysis was performed on Fiji using a script directly inspired by the FRAP Tools script of The Hardin Lab (https://worms.biology.wisc.edu/research/microscopy/), with the following modifications: frame 2 as bleaching frame instead of the frame with minimum intensity, and the first channel (AF488) as the one to be analyzed for fluorescence intensities. Briefly, this Fiji/Python script measures fluorescence intensities in a bleached region of interest (ROI; a random bacterial pole) and a reference ROI (the opposite pole). The data are double normalized using the pre-bleach frame (frame 1) and bleach frame (frame 2) to correct for acquisition bleaching and intensity fluctuations. Recovery data were exported into GraphPad Prism10 and plotted. The script is available in **Suppl. data files**.

Note that the same labeling protocol was used on wild-type bacteria in **Suppl. Fig. 3**, but with Texas Red Succinimidyl Ester (TRSE, Invitrogen) at 1 μg mL^−1^ instead of AlexaFluor™ 488 NHS Ester (Invitrogen). Data were visualized using classical epifluorescence microscopy as described above.

### Thin layer chromatography (TLC)

Lipids were extracted using a modified Bligh and Dyer protocol at a 1:20 (v/v) sample-to-solvent ratio as previously described^74^. Bacterial pellets were weighed and stored at −80 °C or processed immediately. Dry pellet mass was determined by weighing, and samples were resuspended in water to obtain an 80% (w/v) aqueous suspension before transfer to glass tubes. All subsequent steps were performed in glassware to avoid solvent interaction with plastics. Lipids were extracted by adding 350 µL chloroform and 700 µL ice-cold methanol per 100 µL, followed by vortexing and incubation on ice. A second addition of chloroform was performed, and samples were incubated for 30 min on ice with intermittent vortexing, followed by addition of 630 µL _dd_H_2_O and a further 10 min incubation on ice. Phase separation was achieved by centrifugation for 10 min at 2,000 g and 4°C, and the lower organic phase was recovered using glass pipettes. The aqueous phase was re-extracted with chloroform/methanol (1:1), centrifuged under the same conditions, and the organic phases were pooled. Extracts were dried under an inert gas stream and resuspended in chloroform/methanol/acetic acid/_dd_H_2_O (100:20:12:5). Samples were stored in airtight glass tubes with Teflon caps under inert atmosphere at -20°C and protected from oxygen exposure. Lipid extracts were analyzed by thin-layer chromatography (TLC) on silica gel 60 plates using chloroform/methanol/acetic acid/ _dd_H_2_O (100:20:12:5) as the mobile phase. Glycolipids were visualized by spraying with orcinol reagent (200 mg orcinol in 11.4 mL H₂SO₄, brought to 100 mL with _dd_H_2_O) followed by heating at 110°C for approximately 5 min until band development.

### Sacculi analysis

Exponentially growing bacteria (OD_600_ ∼ 0.5-1.5) were harvested by centrifugation and sacculi were extracted by boiling in 4% SDS for 45 min. Sacculi were collected by centrifugation at 100,000 x g for 20 min at 20°C and washed five times with H_2_O. Sacculi were treated with 1 µL of proteinase K (20 mg mL^−1^) in 1 mL of 100 mM Tris-HCl (pH 8) for 1h at 37°C, then the reaction was stopped by adding 110 µL of SDS 10% and boiling for 5 min. Pellets were washed five times with H_2_O at 100,000 x g for 20 min to remove SDS. Purified sacculi were treated with 100 µg mL^−1^ trypsin in 20 mM Tris-HCl (pH 7.5) for 16 h at 37°C, washed four times with H_2_O, incubated for 10 min at 98°C to inactivate the remaining enzyme, and washed a final time with H_2_O. Purified sacculi were treated with 200 µg mL^−1^ α-amylase in 20 mM Tris-HCl (pH 7.5) for 16 h at 37°C, washed four times with H_2_O, incubated for 10 min at 98°C to inactivate the remaining enzyme, and washed a final time with H_2_O. Sacculi were digested with mutanolysine (125 U/mL) in 50 mM ammonium bicarbonate (pH 7.4) with shaking at 1000 rpm for 16 h at 37°C. Insoluble debris was removed by centrifugation, and the soluble fraction containing the muropeptides was reduced with sodium borohydride at 5 mg mL^−1^ in 125 mM borate buffer (pH 9.0) for 45 min. The reaction was acidified to pH 3.0 with a 20% formic acid solution, lyophilized, and resuspended in mobile phase A (H_2_O, 0.1% formic acid). Muropeptides were separated by injecting 5 µL of sample on a C18 column (Hypersil GOLD aQ, 150 x 2.1 mm, 1.9 µm; Thermo Fisher Scientific) at 50°C on an Agilent 1290 HPLC system. The flow rate was constant at 0.4 mL/min using mobile phase A and B (acetonitrile, 0.1% formic acid). An Agilent 6546 ion funnel mass spectrometer (MS) was used in the positive mode with an electrospray ionization (ESI) (voltage 3500 V, Nozzle voltage 1000 V, sheath gas 350°C at 11 L/min, nebulizer pressure 35 psi and drying gas 300°C at 8 L/min). Elution was performed with a linear gradient (0 to 20%) of mobile phase B applied between 10 and 80 min.

### Tn-seq experiment

A transposon sequencing (Tn-seq) library was generated by conjugation between *S. yanoikuyae* and *E. coli* MFDpir carrying the mini-Tn5 transposon encoded on the pXMCS-2 (KanR) plasmid^75^. Overnight cultures were grown in TSB at 30°C for *S. yanoikuyae* and in LB supplemented with DAP and kanamycin at 37°C for *E. coli*, then subcultured and incubated at 37°C to mid-exponential phase. Cultures were mixed (1:1), pelleted by centrifugation, washed twice with TSB, and concentrated prior to mating by spotting 100 µL drops on TSB plates supplemented with DAP and incubating overnight at room temperature. Conjugation mixtures were recovered with a loop and resuspended harshly by pipetting into liquid TSB. Bacteria were allowed to sediment for 1 minute and non-sedimented bacteria were spread on TSB plates supplemented with 30 μg mL^−1^ kanamycin and 50 μg mL^−1^ streptomycin, then incubated for 48h at 30°C. Bacteria were recovered in Recovery Buffer (250 mM EDTA, 10% PEG8000, pH 8) and pelleted at 14 000 G for 20 min at 4°C. Genomic DNA was extracted using a two-step procedure involving manual removal of exopolysaccharides followed by purification with the NucleoBond AXG500 kit (Macherey-Nagel). The library preparation and sequencing were done by Fasteris. Essentiality analysis was performed using TnBox (https://github.com/fxstubbe/TnBox), a combined pipeline described before^39^. Briefly, we used a sliding-window insertion profiling (R100S10; 100 bp windows shifted by 10 bp, scoring insertion-free regions) and transposon insertion frequency (TnIF), calculated as log-transformed normalized read counts per nucleotide position divided by gene length. Genes were classified as essential when at least one 100 bp window lacked insertions and TnIF values were below two standard deviations from the mean, while genes with low fitness or undetermined status were defined based on TnIF thresholds and window-based insertion patterns. Detailed data for **Fig. 6A** are available in **Suppl. Table 1**. Detailed data for the rest of the genome are available upon request.

## Supporting information

Suppl. Data (FRAP script)

## Data Availability

The data supporting this study are available upon reasonable request from the corresponding author.

## Acknowledgments

We thank members of the URBM for stimulating discussions, and more particularly for their help in plating the transposon bank. We thank X. De Bolle for his trust and financial support. We thank the MorphIm platform (https://www.unamur.be/en/technology-platform/morph-im), and more particularly H.F. Renard and B. Ledoux for their help with FRAP experiments imaging. We thank UNamur (https://www.unamur.be/) for financial and logistic support. H.V. was supported by an EMBO postdoctoral fellowship (ALTF 359-2023) and a Postdoctoral Researcher (CR) fellowship from FRS-FNRS; N.Y.M. was supported by Fundação para a Ciência e a Tecnologia (FCT) through Individual Call to Scientific Employment Stimulus (2022.00548.CEECIND) and Call for Exploratory Projects in All Scientific Domains 2023 (2023.12822.PEX); L.W. and J.F.C. were supported by a WELBIO grant from the Walloon Region; CMMI is supported by ERDF funding and the Walloon Region; K.P. was supported by a Postdoctoral Researcher (CR) fellowship and a Research Associate (CQ) mandate from FRS-FNRS.

## Author Contributions

Conceptualization, methodology and data curation were done by K.P. Experiments were performed by S.C., H.V. under the supervision of J.F.C., L.W, E.H. and K.P. Data analysis was performed by S.C., H.V., N.Y.M. and K.P. The manuscript was written by K.P.

## Competing interests

The authors declare no competing interests.

## Notes

### Competing Interest Statement

The authors have declared no competing interest.

### Summary of Updates

- New Suppl. Fig 4: analysis between fluorescence of septum vs membrane HADA signal - Updated Fig 3: graph from n=3 instead of representative experiment - Updated Fig 4: better images for rf470DL - Updated Fig 5: n=3 for PG analysis (compared to n=2 before)

