## Supplementary material for "Extreme envelope plasticity drives temperature-dependent morphogenesis in the LPS-free bacterium *Sphingobium yanoikuyae*": Suppl. Figures

**Suppl. Fig. 1**

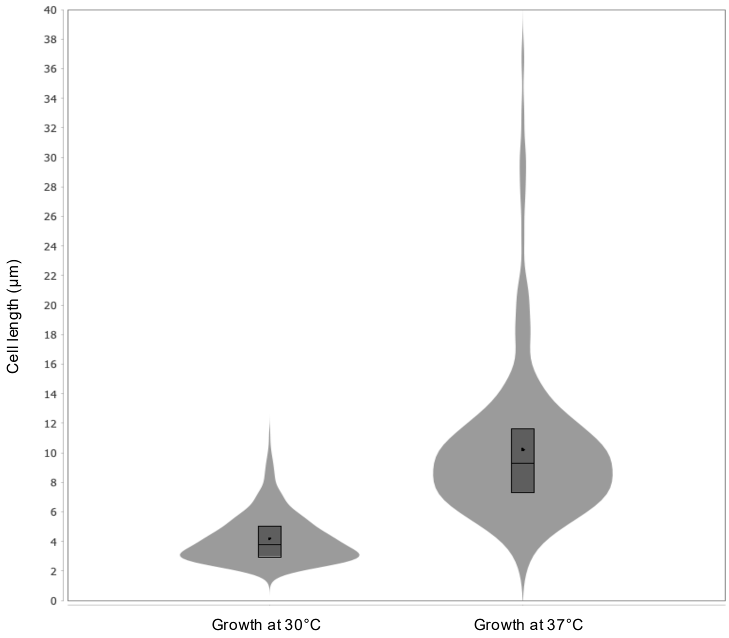


**Suppl. Fig. 1. Violin plot of cell lengths depending on growth temperature.**
The box represent the interquartile range, *i.e.* the middle 50% of the data. The median value is represented as a black line and the mean value as a black dot. n = 479 and 209 cells, respectively.

**Suppl. Fig. 2**

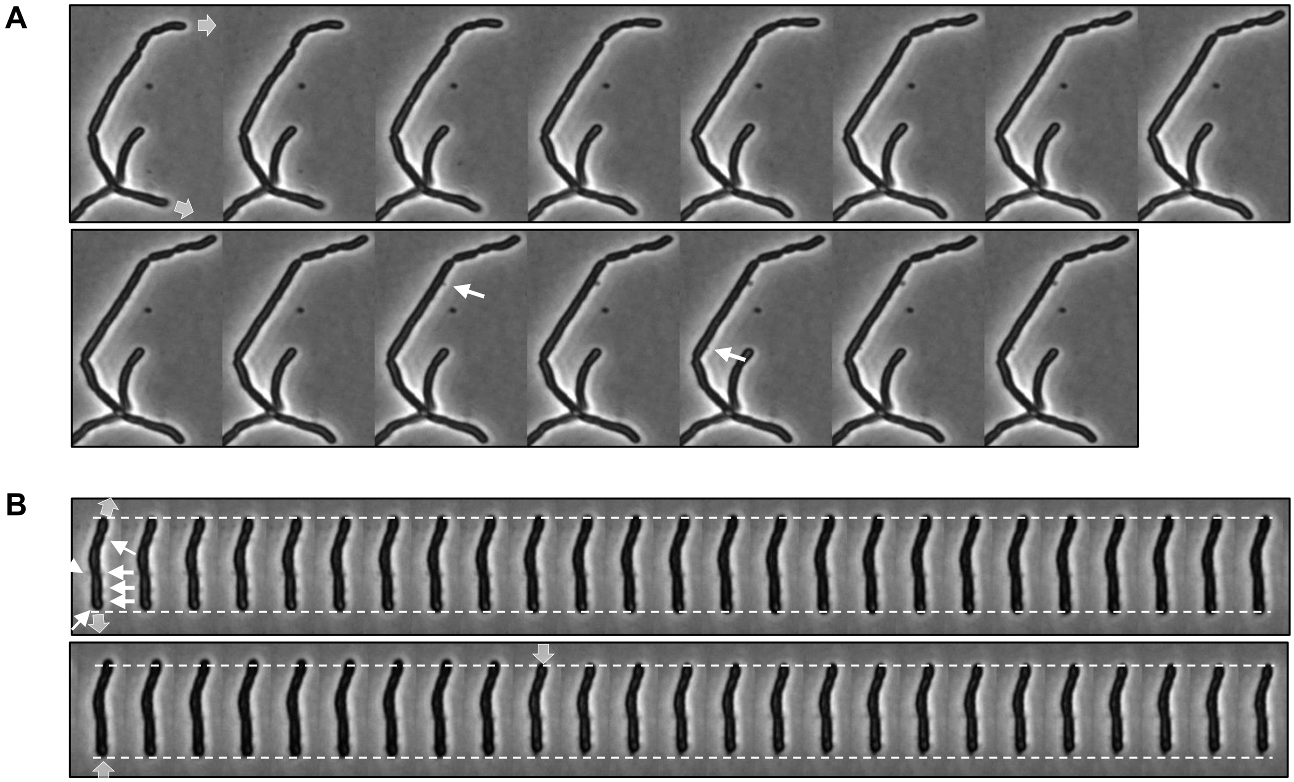


**Suppl. Fig. 2. Time lapse experiments on TSB agar pads at 37°C.**
**A.** Example of a bacterium growing on pad (grey arrows). Images were taken every 5 minutes. After 30 minutes (frame 7), growth was halted and outer membrane vesicles (OMVs) started to appear (white arrows). **B.** Another example of a bacterium slightly growing at 37°C. Images were taken every 5 minutes. Dotted lines are static and help visualize the growth evolution over time. Gray arrows point in the direction of growth (outward) and "retractation" (pointing inward), potentially due to the high amount of OMVs (white arrows) produced.

**Suppl. Fig. 3**

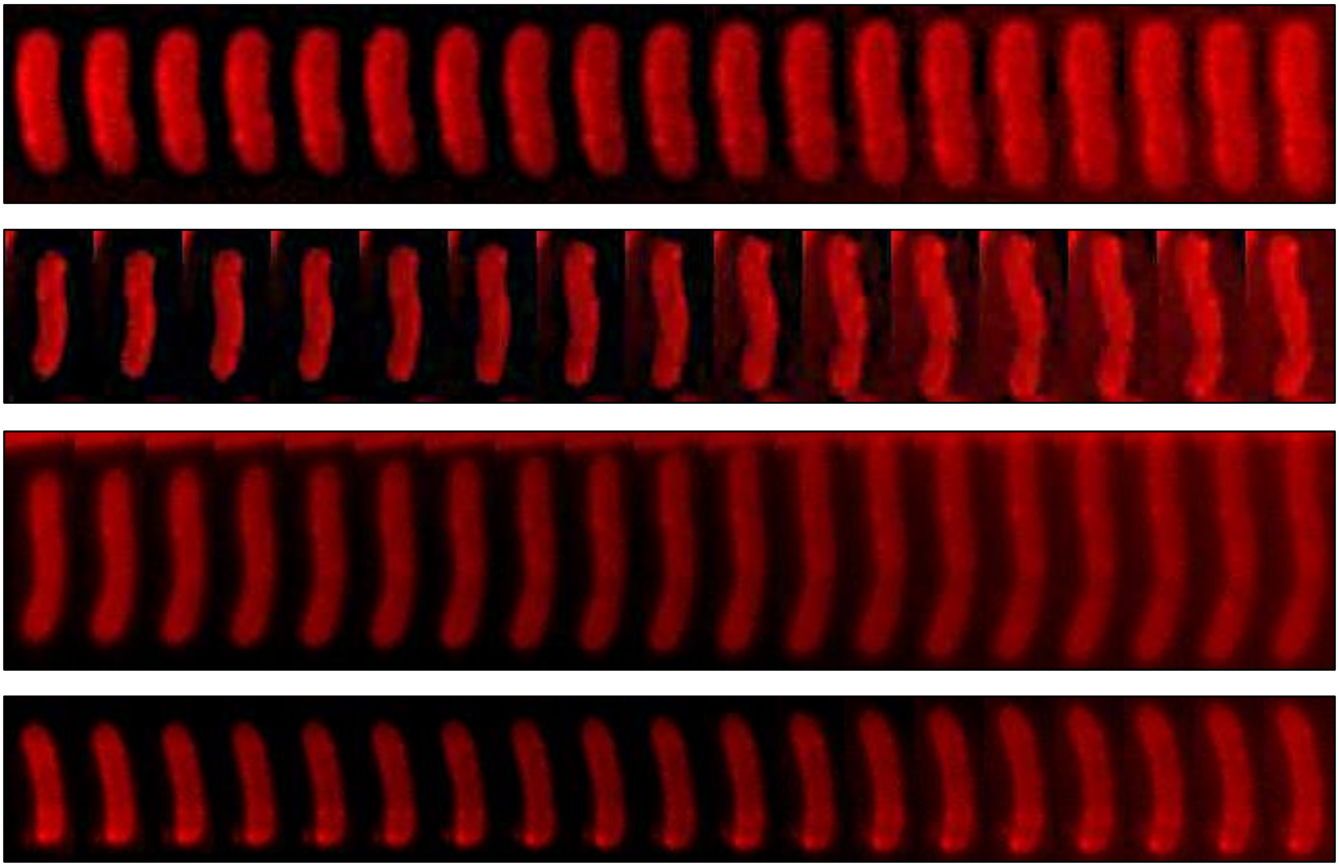

**Suppl. Fig. 3. Time lapse experiments of TRSE-labeled *S. yanoikuyae* on TSB agar pads at 30°C.** Growth pattern of four representative cells pulse-labeled with Texas Red Succinimidyl Ester (TRSE). Images were taken every 5 minutes. See Brown et al, 2012 for comparison with unipolar growing *A. tumefaciens* TRSE labeling, for example.

**Suppl. Fig. 4**

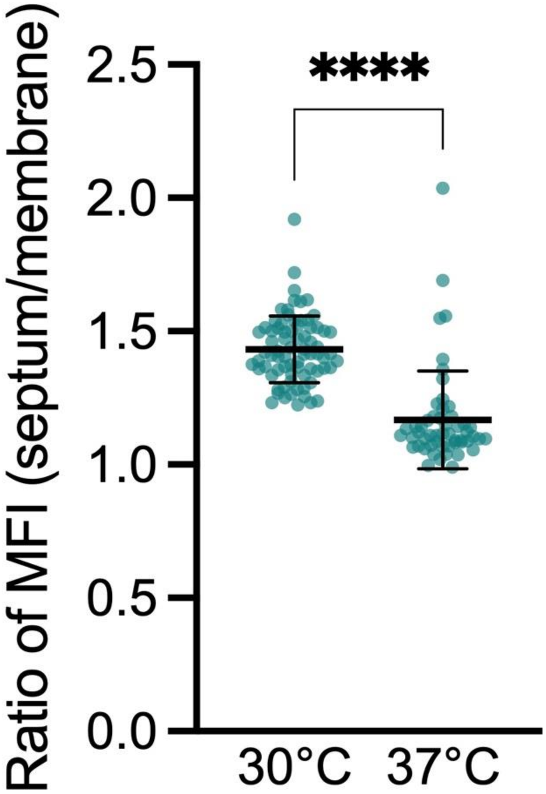


**Suppl. Fig. 4. Analysis of HADA labeling.**
The activity of periplasmic transpeptidases at the septum and on lateral membranes was assessed through HADA labeling of the peptidoglycan. Bacteria were washed and fixed after 5 minutes of incubation with HADA, then visualized by microscopy. Ratio of mean fluorescence intensities (MFI) between septum and membrane were calculated for pre-divisional bacteria grown at 30°C or 37°C (n = 75 and 53 ratio, respectively). Statistical analysis using unpaired Welch's *t* test revealed a highly significant difference between the two conditions (****, $p$ value < 0.0001).
